# Single cell genotypic and phenotypic analysis of measurable residual disease in acute myeloid leukemia

**DOI:** 10.1101/2022.09.20.508786

**Authors:** Troy M. Robinson, Robert L. Bowman, Sonali Persaud, Ying Liu, Qi Gao, Jingping Zhang, Xiaotian Sun, Linde A. Miles, Sheng F. Cai, Adam Sciambi, Aaron Llanso, Christopher Famulare, Aaron Goldberg, Ahmet Dogan, Mikhail Roshal, Ross L. Levine, Wenbin Xiao

## Abstract

Measurable residual disease (MRD), defined as the population of cancer cells which persists following therapy, serves as the critical reservoir for disease relapse in acute myeloid leukemia (AML) and other malignancies. Understanding the biology enabling MRD clones to resist therapy is necessary to guide the development of more effective curative treatments. Discriminating between residual leukemic clones, preleukemic clones and normal precursors remains a challenge with current MRD tools. Herein, we developed a single cell (sc) MRD assay by combining flow cytometric enrichment of the targeted precursor/blast population with integrated scDNA sequencing and immunophenotyping. Our scMRD assay shows high sensitivity of approximately 0.01%, deconvolutes clonal architecture and provides clone-specific immunophenotypic data. In summary, our scMRD assay enhances MRD detection and simultaneously illuminates the clonal architecture of clonal hematopoiesis/pre-leukemic and leukemic cells surviving AML therapy.

**Statement of significance:** ScMRD assay integrates mutation and immunophenotype at single cell resolution and therefore distinguishes clonal hematopoiesis/preleukemic vs. leukemic clones. This study serves as a framework for identifying high-risk MRD clones and improving our understanding of both the molecular drivers and vulnerabilities of therapy resistant AML clones.

## Introduction

Acute myeloid leukemia (AML) is a heterogeneous set of hematologic malignancies, characterized by expansion of immature myeloid blasts^1^. Although most patients with AML show an initial response to therapy (60-80%), relapse remains the fundamental challenge to achieving durable cures^1^. Measurable residual disease (MRD) represents a critical, therapy-resistant cancer cell reservoir responsible for disease recurrence^2^. Accurately identifying relevant MRD clones is necessary to risk stratify patients and to guide further therapy to prevent overt relapse and to achieve durable remissions.

MRD is defined by the presence of residual leukemic cells which are not detectable by conventional morphologic assessment. MRD status has significant prognostic value in AML independent of pre-treatment genetics/cytogenetics^2^ and the presence of MRD is associated with adverse outcomes in a spectrum of AML therapies – including chemotherapy, venetoclax-based therapies, and allogeneic stem cell transplant^3-6^. The most common MRD detection methods use multi-color flow cytometry (MFC) analysis of blasts or real-time quantitative polymerase chain reaction (RT-qPCR) of specific mutations or gene fusions detected at diagnosis. MFC MRD-testing relies on abnormal immunophenotypic profiles which may not be readily identifiable in all leukemic cells. The current sensitivity of MFC MRD-testing is 0.1%, as defined by European Leukemia Net (ELN) MRD guidelines^2^. In addition, flow cytometry is frequently unable to distinguish between phenotypic abnormalities of regenerating precursors, clonal hematopoiesis (CH), and residual leukemic blasts, and therefore cannot fully predict the leukemic potential of an immunophenotypically abnormal population. RT-qPCR is highly sensitive and specific, however the utility of this method is limited as only a small percentage of AML patients have a detectable, leukemia-specific mutation or gene fusion that is informative for molecular tracking.^7^ Moreover, it is not feasible to simultaneously detect/quantitate the spectrum of mutations specific to resistant AML clones.

Recently, bulk-tumor next generation sequencing (NGS) has emerged as a promising technology to expand on the arsenal of MRD monitoring techniques^7^. While the sensitivity of bulk NGS (including error corrected) and MFC-based MRD testing may be further improved, the specificity of these analyses faces both theoretical and empirical barriers due to the clonal complexity of AML^8^.

Despite the utility of these assays, there remains a pressing clinical need for discrimination of residual leukemic cells from the mutated CH/pre-leukemic cells that do not invariably portend relapse^6,9^. Residual subclones may have different leukemic potential while ancestral clones may only result in CH rather than frank AML^10^, which cannot be accurately delineated by bulk MRD assays. In addition, bulk NGS MRD assays may detect mutations in mature populations lacking leukemic potential, which is often seen in patients receiving differentiation inducing therapies (e.g. IDH1/2 inhibitors)^11^. These biological complexities require a nuanced interpretation of MRD assays, ultimately hampering the clinical utility of bulk-sequencing and MFC based approaches. With 10-30% of MRD-negative and 40-70% of MRD-positive patients ultimately relapsing^12^, there remains a critical need for a sensitive and specific MRD assay to inform clinical intervention.

Single cell (sc) DNA sequencing technology has recently been utilized to study clonal architecture in AML, distinguish CH/pre-leukemic versus leukemic clones, and detect mutations in remission samples^8,13-15^. To build upon these pioneer studies and resolve the challenges associated with both bulk NGS and MFC MRD-testing, we have developed a novel scMRD assay by combining flow cytometric enrichment of the targeted precursor/blast population with integrated scDNA sequencing and immunophenotyping.

## Results

We hypothesized that combining precursor/blast enrichment and scDNA+protein sequencing technology would increase the sensitivity of MRD detection and delineate MRD clonal architecture. We reasoned that the MRD clones responsible for relapse primarily reside in immature compartments, thus we utilized flow cytometry assisted cell sorting (FACS) to enrich viable CD34^+^ and/or CD117^+^ progenitors prior to loading cells onto the Mission Bio Tapestri single cell sequencing platform (**Fig 1A**). The custom scDNA panel contains 109 amplicons covering 31 genes known to be involved in hematologic malignancies^8^. To increase assay throughput, we multiplexed samples from different patients into each integrated scDNA+protein run. The results of the multiplexed runs were then computationally deconvoluted and used for both single-sample and aggregated analyses.

**Figure 1.**
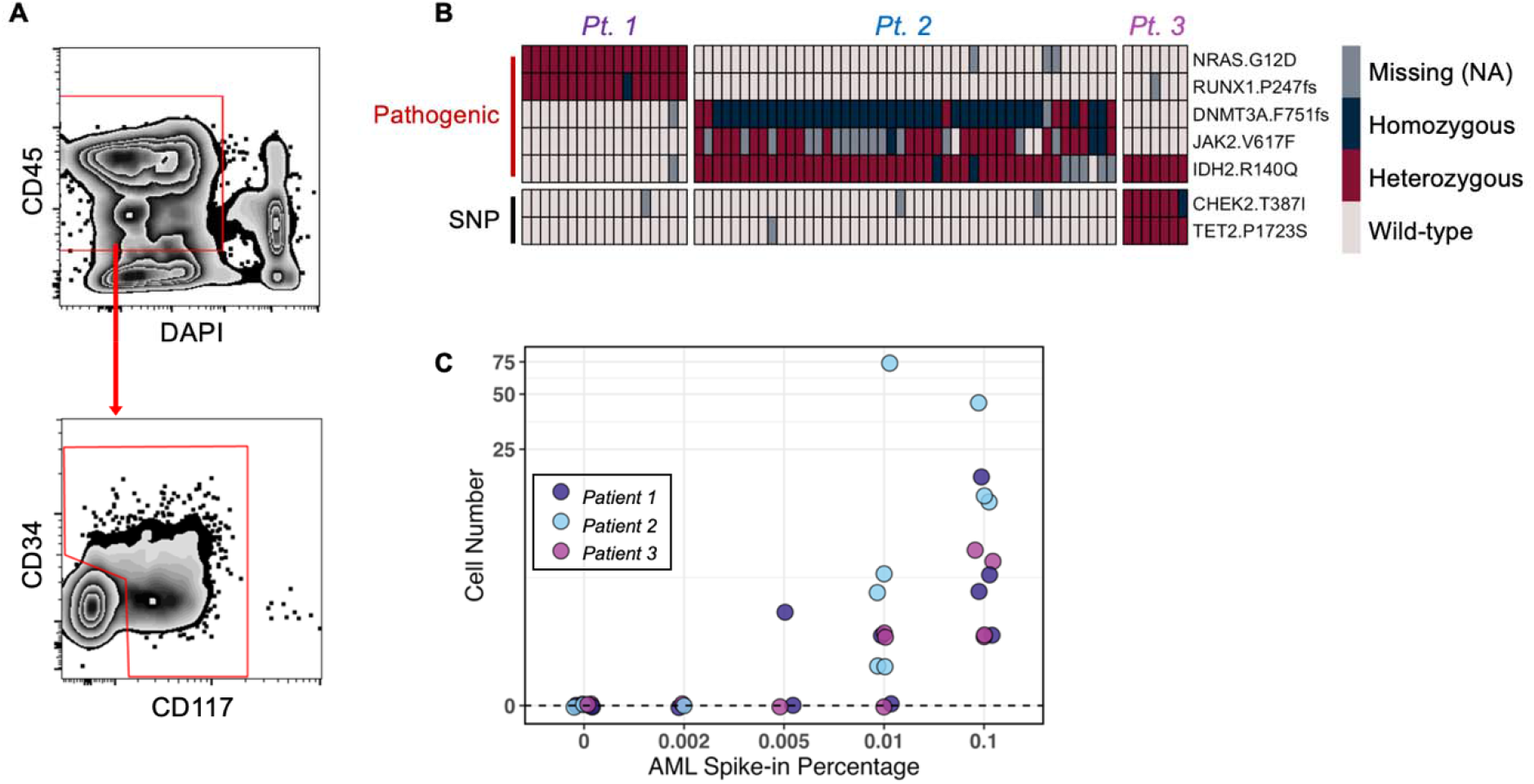
Limit of mutation detection with the scMRD assay. **A**. Schema of gating strategy for flow cytometric enrichment of live CD34^+^ and/or CD117^+^ cells were sorted. For clinical samples, the abnormal blasts were positive for CD34 and/or CD117. **B**. Representative heatmap showing mutation calling of spiked-in AML blasts in a limit of detection experiment testing a sensitivity of 0.1%. **C**. Summary of mutation detection at various sensitivity levels. This plot represents two independent experiments.

To evaluate the sensitivity of the scMRD assay, we performed a limiting dilution study. AML blasts from 3 genetically distinct AML samples harboring clonal mutations were mixed with 10 million normal bone marrow mononuclear cells to test different sensitivity thresholds (10,000 cells 0.1%, 1,000 cells 0.01%, 500 cells 0.005% and 200 cells 0.002%). Mutations were as follows: Patient 1: *NRAS* p.G12D/*RUNX1* p.P247fs, Patient 2: *JAK2* p.V617F/*IDH2* p.R140Q/*DNMT3A* p.F751fs, Patient 3: *IDH2* p.R140Q). FACS-enriched CD34^+^ and/or CD117^+^ cells were multiplexed, subjected to the Tapestri v2 microfluidics platform, and sequenced. Expected pathologic mutations were identified in all 11 replicates at a sensitivity of 0.1% (**Fig. 1B**). Mutations were also identified in 8/10 and 1/3 replicates at a threshold of 0.01% and 0.005%, respectively, and mutations were not identified in blank controls (0/9 replicates) or when present at 0.002% (0/3 replicates) (**Fig. 1C**). Limiting dilution analysis estimated a sensitivity of 0.0077% (95% CI [0.004% – 0.0153%])^16^. These data demonstrate the high sensitivity and specificity of mutation detection using the scMRD assay.

We next applied our scMRD assay to 30 cryopreserved post-induction chemotherapy MRD samples obtained from 29 AML patients (median age 71 years old, 15 male and 14 female, see **Supplementary Table S1**). MRD was scored as negative in 2 samples by MFC and in 6 samples by bulk NGS (**Fig 2A**). The median cell number of these samples was 2.6 million (ranging from 0.6-14.1 million) with a viability range of 27-55%. FACS-enriched CD34^+^ and/or CD117^+^ viable cells were multiplexed with up to 5 unique patient samples per run and processed via the Tapestri platform (50-100 thousand cells per run, median 65 thousand) (**Supplementary Fig S1A**).

**Figure 2.**
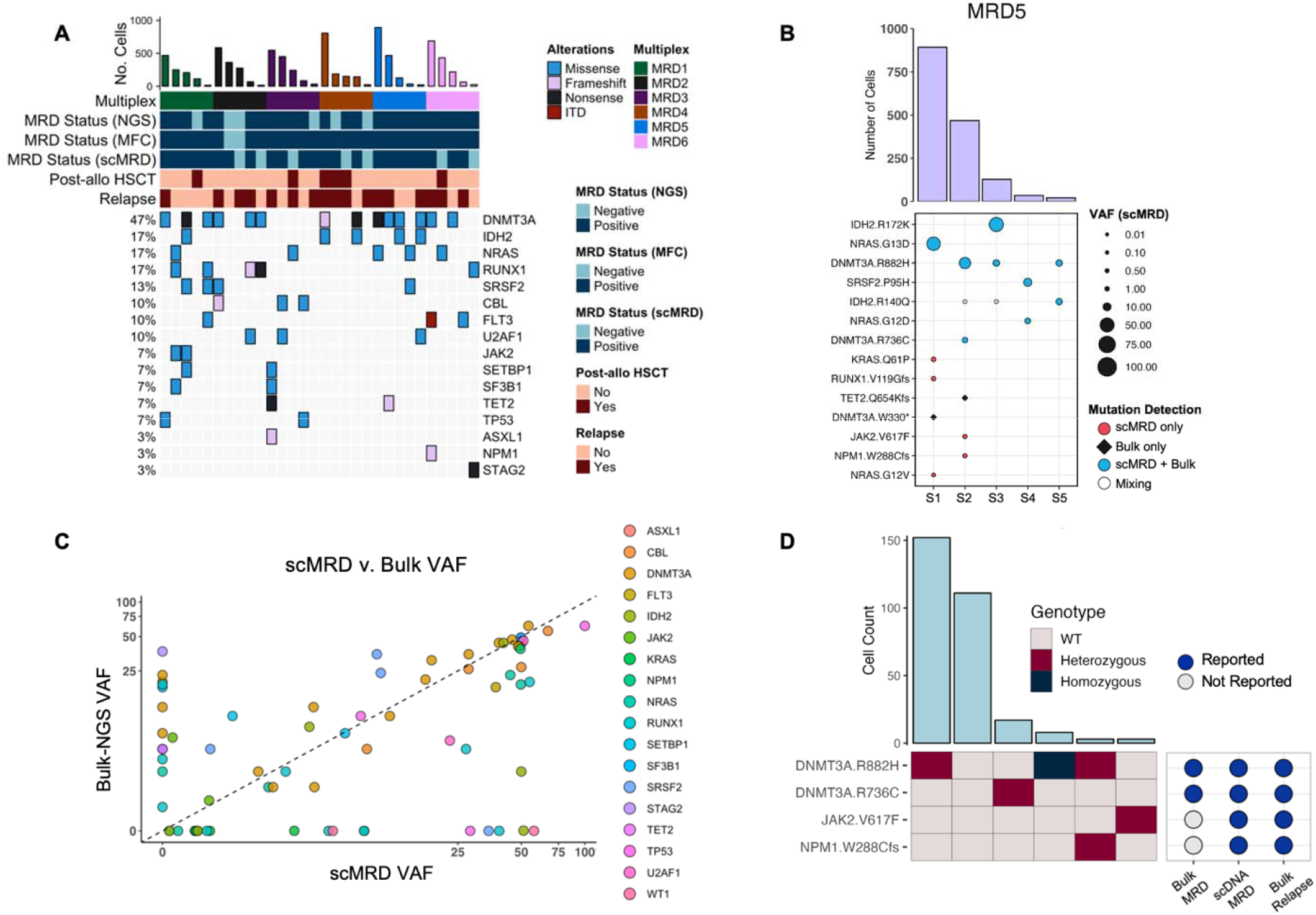
Mutation and relapse associated clone identified by scMRD assay. **A**. Oncoprint showing concordance of MRD detection by bulk NGS assay, scMRD assay and MFC. Bar plot (top) represents the number of cells recovered after computational demultiplexing. Mutations represent those that were detected by bulk NGS at the remission timepoint and are covered by the custom scDNA panel. Post-allo HSCT represents the time of MRD assessment. Relapse represents outcomes after MRD assessment. **B**. A representative deconvolution plot of one multiplexed scMRD run. **C**. Comparison of mutations detected by bulk NGS vs scMRD. **D**. Clonograph of a patient (MRD5-S2) illustrating scMRD-specific detection of *NPM1* and *JAK2* mutations that were present at late relapse.

Given that different samples from multiple patients were included in each scMRD run, we developed a computational approach to deconvolute different individuals in each sequencing run at the single cell level. Downstream demultiplexing of scMRD sequencing data used germline single nucleotide polymorphisms (SNPs) covered by the custom scDNA panel (**Supplementary Fig S1, Supplementary methods**). We filtered our dataset based on genotyping call rate to include cells with complete genotyping information for the top 10-20 SNPs in each run and performed K-means clustering on cells with non-missing SNP allele frequencies. To identify and remove doublets within each scMRD run, we implemented a method to simulate artificial doublet SNP profiles and exclude putative real doublets based on similarity to an artificial cluster center (**Supplementary Fig S1, Supplementary methods**). We first randomly sampled our dataset to produce a pool of cells with even representation of each cluster. From this cell pool, we sampled two cells at a time, averaged their SNP allele frequency profiles, and re-clustered the artificial doublets with real cells. We then applied a Euclidean distance metric to assess the similarity between the SNP profiles of real cells and artificial cluster centers. After doublet detection and exclusion, we then classified additional cells in our dataset according to their germline SNP profile. To achieve this, we calculated a Hamming distance between each cell and the most common SNP profile for each cluster. Cells were assigned to clusters based on the SNP profile matching at least 80% of one cluster while being the maximum Hamming distance from every other cluster. This approach enabled us to deconvolute multiplexed scMRD runs and assign sequenced cells to the specific patient from which they were derived without the need to leverage patient-specific somatic mutation information. Using this approach, we classified an average of 1,333 sorted cells per run (1,053 – 1,544 cells, SD = 163.4). Importantly, demultiplexing enabled assignment of hotspot mutations (i.e., *DNMT3A* p.R882H) present in multiple samples within the same multiplex (**Fig 2B** and **Supplementary Fig S1-S2**).

Overall, the results for MRD status and mutation presence were concordant between bulk NGS and scMRD in 22/30 (73%) samples and for 46/77 (60%) mutations, respectively (**Fig 2A**). Mean variant allele frequencies (VAF) of mutations detected by both scMRD and bulk NGS trended towards higher allele burden by scMRD (p=0.064, paired Wilcoxon signed rank test) (**Fig 2C**). Among the 31 discordant mutations covered by both scMRD and bulk NGS panels, scMRD identified 17 mutations that were missed/unreported by bulk NGS, including *RUNX1* (n=5), *NPM1* (n=3), *KRAS* (n=2), *IDH2* (n=2), *WT1* (n=2), *JAK2, TP53* and *SRSF2* mutations (**Supplementary Fig S2)**, 14 (82.4%) of which were associated with and present at relapse. Conversely, there were 14 mutations that were detected by bulk NGS but missed by scMRD, including *DNMT3A* (n=4), *RUNX1* (n=3), *NRAS* (n=2), *STAG2, TET2, SETBP1, SRSF2* and *FLT3TKD* mutations. Interestingly, only 6/14 (42.8% vs 82.4%, p=0.03, Fisher exact test) were present at relapse. There were 4 MRD samples with 7 mutations (2 *RUNX1*, 2 *DNMT3A*, 2 *NRAS*, and *STAG2*) not detected by scMRD. Although these samples had slightly lower viable and recovered cell numbers compared to others (median: 1.5 million vs 2.6 million [p=0.08], 64 vs 175 [p=0.1], respectively, Mann-Whitney test), the cause may be multifactorial including that the presence of specific mutations (i.e. *NRAS*) may reside in mature/differentiated compartments not sampled with our scMRD assay^8^.

We next assessed the ability of scMRD assay profiling to differentiate MRD based on clonal architecture, including discrimination between single mutant CH/pre-leukemic and leukemic clones. We found that scMRD readily deconvolved CH/pre-leukemic vs. leukemic clonal architecture in demultiplexed samples (**Fig 2D** and **Supplementary Fig S3**). In sample MRD5-S2, bulk NGS detected *DNMT3A* p.R882H, *DNMT3A* p.R736C, and *TET2* p.Q654Kfs (not covered by scMRD panel) mutations at the remission timepoint, while scMRD detected both *DNMT3A* mutations in distinct clones, with one subclone harboring *DNMT3A* p.R882H/*NPM1* p.W288Cfs (*NPM1* scVAF = 1.22%) co-occuring mutations and another with a *JAK2* p.V617F (*JAK2* scVAF = 0.23%) mutation. Importantly, bulk NGS at the time of subsequent relapse revealed the presence of both the *NPM1* p.W288Cfs (VAF = 5%) and *JAK2* p.V617F (VAF = 2%) mutations (**Fig 2D**). These data demonstrate that scMRD enables resolution of residual pre-leukemic clones (i.e. *DNMT3A* alone) and leukemic clones (co-mutant *DNMT3A*/*NPM1*) that persist at relapse.

Integration of scMRD immunophenotypic analysis enabled identification of mutation and clone-specific expression of key cell surface proteins (**Fig 3A-B and Supplementary Fig S4A-B**). Compared to wild type clones, single mutant clones displayed differential expression of CD34, such as *U2AF1* (log2FC = 3.5, *P <* 0.002) and *KRAS* (log2FC = -5.72, *P <* 0.02). Interestingly, *NRAS-*mutant clones had a marked increase in expression of CD33 (log2FC = 1.47, *P* < 1.19×10^−6^) but not CD34 (log2FC = 0.20, *P* < 6.16×10^−10^) consistent with a previous study^10^. More importantly, differential immunophenotypic states were identified when comparing CH/pre-leukemic and leukemic clones within and between patients (**Fig 3B-D, Supplementary Fig 4B-C**). Compared to compound mutant *DNMT3A/NPM1* (with or without *FLT3*) clones, single mutant *NPM1* clones showed reduced CD34 (log2FC = -2.75, *P* < 0.0049) and increased CD117 (log2FC = 2.68, *P* <0.043) surface expression. In addition, *NPM1* clones displayed higher CD33 expression (log2FC = 1.03, *P* <0.04) compared to *DNMT3A* clones. Importantly, this distinct immunophenotype is well described in patients with *NPM1*-mutated AML^14^. We observed highly similar immunophenotypes between wild type and *DNMT3A*-mutant cells, suggesting CH clones (at least with *DNMT3A* mutation) may not have overtly aberrant surface protein expression (**Fig 3C-D**). By contrast, co-mutant *DNMT3A*/*IDH2* or *DNMT3A*/*NPM1* cells showed consistently aberrant immunophenotypes, with *DNMT3A/IDH2* cells characterized by increased CD71 expression (log2FC = 0.388, *P*<8.7×10^−8^, vs. *DNMT3A*) (**Fig. 3C-D and Supplementary Fig 4B-C**), indicative of erythroid biased differentiation consistent with previous reports^17,18^. Concordant with the findings in **Fig 3B**, the overall patterns of surface protein expression were different between *NPM1*-mutated and *DNMT3A/NPM1* co-mutated cells with the latter more highly expressing CD34 and granulocytic/monocytic markers such as CD16 (log2FC = 3.37, *P*<1.2×10^−9^, Percent Expressing = 98.9%) and CD64 (log2FC = 3.05, *P*<7.7×10^−4^, Percent Expressing = 75.9%) (**Fig 3D**). These data highlight that integrated genomic/immunophenotypic analysis at the MRD time point can distinguish between CH/pre-leukemic and leukemic clones that portend a substantively higher likelihood of relapse.

**Figure 3.**
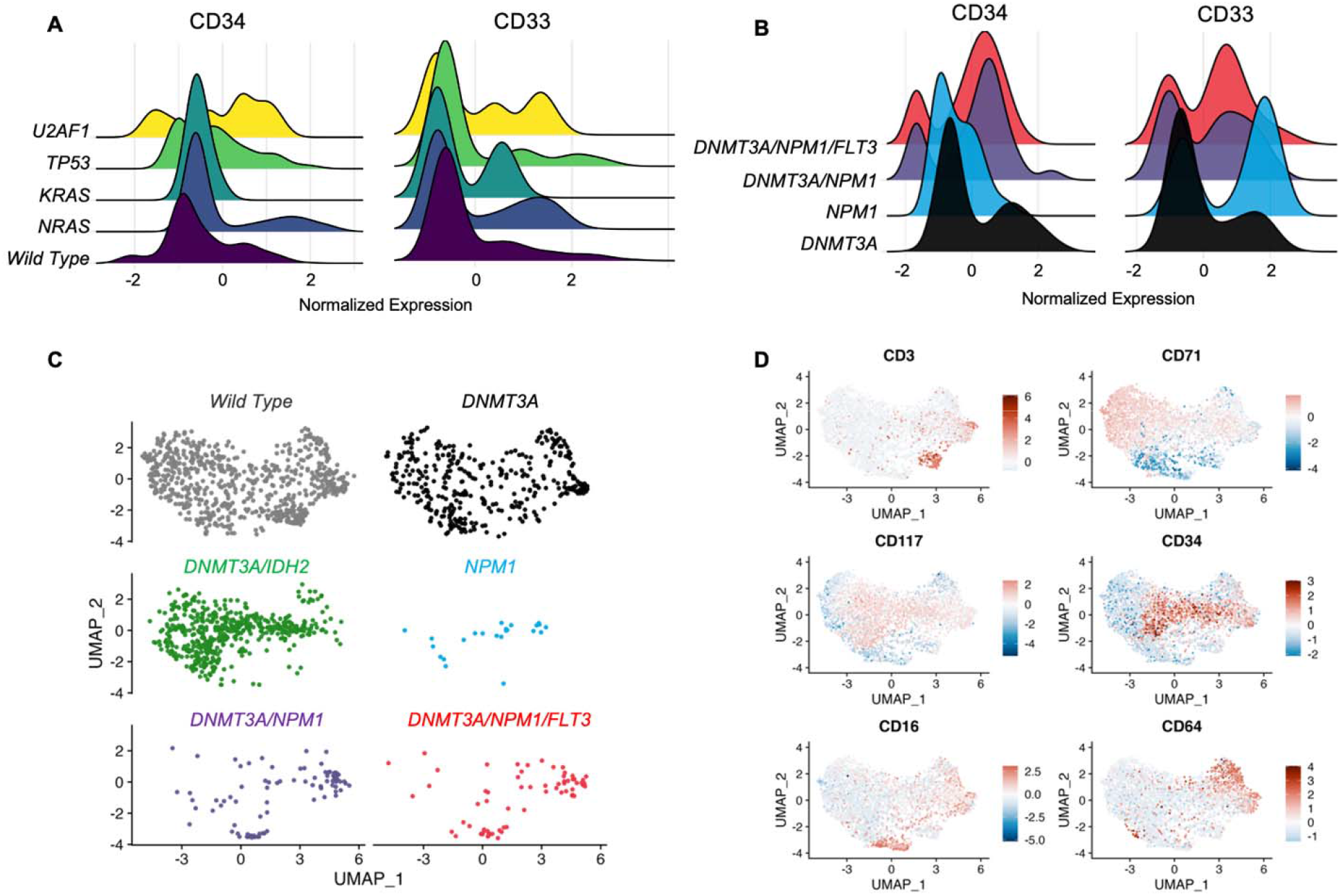
Clone- and mutation-specific immunophenotype. **A**. Clone specific immunophenotype. **B**. Differential surface marker expression between CH/preleukemic vs leukemic clones. **C-D**. UMAP analysis of immunophenotypes of CH/preleukemic vs leukemic clones. Data are log-normalized, centered, and scaled on a by-run basis.

Within our cohort, 5 samples were obtained post-allogeneic hematopoietic stem cell transplant (HSCT) and represented an admixture of donor and recipient cells. We therefore asked if scDNA sequencing can distinguish host and donor derived cells post-allogeneic HSCT. Germline SNP-based deconvolution identified distinct non-mutant clusters consistent with donor origin, which were confirmed as donor samples by matching the SNP profile of paired pre-transplant samples. Donor-host pairs were successfully recovered for all 5 samples (**Fig. 4A**). Chimerism was calculated based on recovered host vs donor cells and correlated with the results by bulk short tandem repeat genotyping (STR) on unsorted BM samples. The levels of host cells detected by scMRD assay were significantly higher than those by STR testing in 4 relapsed patients (median of difference 30%, p=0.04, Wilcoxon matched pairs signed rank test), suggesting that host cells show enrichment in immature compartments and represent an early indicator of relapse. Integrated immunophenotypic analysis of post-allogeneic HSCT samples showed distinct cell surface protein expression between donor and host cells. Analysis of sample MRD1-S4 revealed clear separation of donor and *NPM1*-mutant host cells, with the former containing a subset of spiked-in CD3^+^CD8^+^ T-cells, and the latter displaying aberrant increased expression of CD33 (log2FC = 2.54, *P*<1.32×10^−4^), CD13 (log2FC = 3.9, *P*<1.06×10^−5^), and CD123 (log2FC = 4.08, *P*<0.011) (**Fig 4B**), of which CD123 is a well described leukemic stem cell marker^19,20^. In addition, expression of the T-cell activation marker, CD69 was observed in a subset of donor and host T cells but also unexpectedly in host leukemic cells, consistent with previous studies showing that CD69 may be expressed in leukemic stem cells and thus may represent a surface marker for MRD detection^21^. Importantly, abnormal immunophenotype of these host leukemic blasts identified by MFC was also detectable by scMRD, with elevated co-expression of CD33 and CD117 on *NPM1*-mutant cells and characteristically low levels of CD34 (**Fig 4C**).

**Figure 4.**
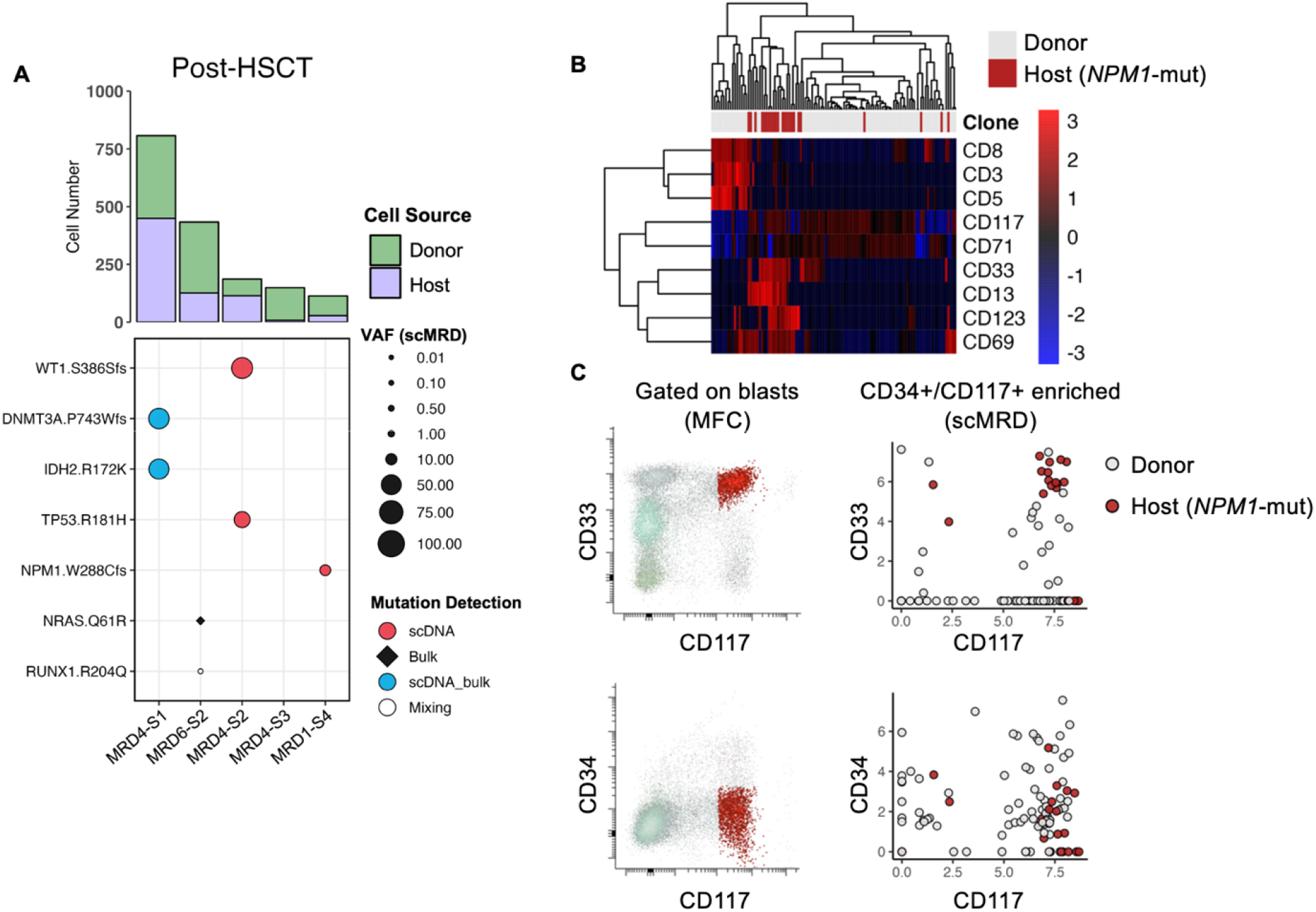
scDNA + protein analysis enables simultaneous identification of donor cells and MRD. **A**. Aggregated deconvolution plot showing mutations detected and host-donor chimerism of post-allogeneic HSCT samples included in the study. MRD4-S3 had an *HDAC1* P243L mutation not covered by the scMRD panel. **B**. Heatmap analysis of differential surface maker expression between donor and host cells in MRD1-S4. **C**. Concordance of immunophenotype of MRD cells between MFC and scMRD in MRD1-S4.

## Discussion

Our study illustrates the feasibility of single cell genotypic and immunophenotypic profiling at the remission timepoint to enumerate and delineate MRD through blast enrichment and scDNA+protein technology. Our data demonstrates that scMRD profiling readily resolves clonal architecture and can distinguish between single mutant CH/pre-leukemic vs. leukemic clones with multiple co-occurring mutations. The integration of mutation and immunophenotypic information further enhances MRD detection by identifying genotype-specific protein expression patterns. This can be potentially utilized to isolate relevant clones for studying MRD biology and therapeutic vulnerabilities. Given the increased use of molecular/cell surface-targeting therapeutic modalities for AML patients^22,23^, assessing expression of surface markers in relevant MRD clones with defined mutational repertoires may provide further guidance for treatment. Future integration of scDNA+protein sequencing with clone-level transcriptional analysis will provide additional insights into the molecular pathways mediating leukemic cell persistence following different AML therapies.

The multiplexing approach and deconvolution algorithm implemented in this study is reliable and cost effective, however, more work is needed to optimize this assay. While scMRD detected many mutations missed by bulk NGS, some mutations were also missed by scMRD. There are several explanations: first, the starting cell numbers of these samples with mutations missed by scMRD assay were slightly lower, leading to lower cell recovery compared to clinical profiling performed on fresh samples. In clinical practice, fresh samples with higher cell numbers should be used for scMRD testing, which will certainly improve cell recovery and sensitivity. Second, our use of FACS for blast enrichment overall can alter mutation allele burden compared to bulk sequencing, with some mutations being enriched in immature HSPC compartments and others enriched in differentiated clonal progeny. For example, previous studies have shown *NRAS* mutated clones express high levels of CD11b, a marker often indicative of more differentiated myeloid cells (neutrophils and monocytes)^8^. Consistent with this notion, a subset of *NRAS* mutations were missed by the scMRD assay. Notably, only a small portion of mutations missed by scMRD were associated with relapse. High-purity depletion of mature compartments, as in our flow enrichment, may theoretically decrease the burden of these mutations, including those CH mutations present in mature myeloid progeny with reduced/absent leukemic potential. In clinical practice, magnetic bead purification of stem/progenitors followed by scMRD assessment will allow for accurate detection and delineation of MRD in a more cost-effective manner and can serve as an adjunct to bulk NGS performed pre-therapy and at relapse.

In conclusion, our scMRD assay not only enables sensitive MRD detection, but also achieves sufficient resolution to characterize the clonal architecture of pre-leukemic/leukemic cells that persist after therapy, which will ultimately increase the specificity of MRD results. Larger studies are needed to comprehensively elucidate the clonal architecture of MRD and study the dynamics of clonal evolution by comparing clonal architecture and immunophenotype at diagnosis and relapse. Our study paves the road for delineating the genetic and phenotypic properties of high-risk MRD clones and better understanding the molecular underpinnings of MRD in AML.

## Methods

### Patient samples

Bone marrow aspirates were received in the clinical lab at MSKCC. After 5 days with all necessary clinical tests being completed, the leftover cells were deemed as medical waste and mononuclear cells were obtained by centrifugation on Ficoll from bone marrow and viably frozen. Uninvolved bone marrow aspirates from patients with stage 1 B-cell lymphoma were used as normal controls. Patient samples underwent high-throughput genetic sequencing with an FDA approved targeted deep sequencing assay of 500 genes (IMPACT-heme) or by an NGS platform panel composed of 49 genes that are recurrently mutated in myeloid disorders (RainDance Technologies ThunderBolts Myeloid Panel). Informed consent was obtained from patients according to protocols approved by the institutional IRBs and in accordance with the Declaration of Helsinki. This study was approved by MSK Institutional Review Board (#12-245, #13-037, #16-1591).

### Cell enrichment

Patient samples were thawed, washed with FACS buffer, and quantified using a Countess cell counter. Cells (0.5–4.0 × 10^6^ viable cells) were then resuspended in cell staining buffer (#420201, BioLegend) and incubated with TruStain FcX, and 1× Tapestri blocking buffer for 15 min on ice. Cells were incubated with anti-human CD4 (clone: OKT4)-APC/Cy7 (dilution 1:30), anti-human CD8 (clone: RPA-T8)-BV711 (dilution 1:30), anti-human CD34 (clone: Qbend)-APC (dilution 1:10), anti-human CD117 (clone: A3C6E2)-PE (dilution 1:75), and anti-human CD45 (clone: Q17A19)-AlexaFlour 488 (dilution 1:30) for 15 minutes on ice. Then TotalSeq™-D Human Heme Oncology Cocktail, V1.0 (# 399906, BioLegend) containing the pool of 45 oligo-conjugated antibodies was added and incubated for an additional 30 minutes on ice. Cells were then washed 3 times with cell staining buffer (#420201, BioLegend) followed by resuspension of the cells in DAPI containing FACS buffer. DAPI negative and CD45 positive viable cells were gated. After exclusion of CD4 and CD8 positive lymphocytes, CD34+/CD117-, CD34+/CD117+ and CD34-/CD117+ populations were combined for sorting using a SH800S Cell Sorter. In MRD1 run, 1000 sorted CD4 vs CD8 positive T-cells from two individual samples were spiked in, respectively.

### Single-cell DNA and protein library preparation and sequencing

Enriched cells were resuspended in Tapestri cell buffer and quantified using a Countess cell counter (Invitrogen). Single cells (1,000–3,000 cells/μl) were encapsulated using a Tapestri microfluidics cartridge and lysed. A forward primer mix (30 µM each) for the antibody tags was added before barcoding. Barcoded samples were then subjected to targeted PCR amplification of a custom 109 amplicons covering 31 genes known to be involved in AML. DNA PCR products were then isolated from individual droplets and purified with Ampure XP beads. The DNA PCR products were then used as a PCR template for library generation as above and repurified using Ampure XP beads. Protein PCR products (supernatant from Ampure XP bead incubation) were incubated with Tapestri pullout oligo (5 μM) at 96□°C for 5 min followed by incubation on ice for 5 min. Protein PCR products were then purified using Streptavidin C1 beads (Invitrogen) and beads were used as a PCR template for the incorporation of i5/i7 Illumina indices followed by purification using Ampure XP beads. All libraries, both DNA and protein, were quantified using an Agilent Bioanalyzer and pooled for sequencing on an Illumina NovaSeq by the MSKCC Integrated Genomics Core.

### Data Processing and Variant Filtering

FASTQ files from scDNA+protein samples were processed via the TapestriV2 pipeline as described previously^8^. This analytics platform trims adaptor sequences, aligns sequencing reads to the hg19 reference genome, and calls cells based on completeness of amplicon sequencing reads for each barcode, and calls variants using GATKv3.7 best practices. After pipeline processing, data for each run were aggregated into H5 files, which were downloaded and read into R using the rhdf5 package. Downstream processing was conducted using custom scripts in R, which will be made available at https://github.com/RobinsonTroy/scMRD. Low quality variants and cells were then excluded based on filtering cutoffs for genotype quality score (<30), read depth (<10) alternate allele frequency (<20%), and presence in <0.1% of cells. The details of scMRD computational demultiplexing and scMRD protein analysis are included in supplementary methods.

### Limit of Detection Study Analysis

For each multiplexed AML spike-in run, the numerical genotype matrix (NGT) was extracted from each respective H5 file in R. Each of the three AMLs harbored >1 pathogenic mutation, except for patient 3 which contained a single *IDH2*.*R140Q* mutation. To increase the confidence in accurate cell calling, two additional heterozygous germline SNPs (*CHEK2*.*T387I* and *TET2*.*P1723S*) were identified as private to patient 3. The curated list of known variants included mutations/SNPs present in patient 1 (*NRAS*.*G12D, RUNX1*.*P247fs*), patient 2 (*DNMT3A*.*F751fs, JAK2*.*V617F, IDH2*.*R140Q*), and patient 3 (*IDH2*.*R140Q, CHEK2*.*T387I, TET2*.*P1723S*). After filtering, all cells were queried for variants included in this list. Cells harboring the expected variants were then filtered based on the requirement that real cells must contain at least two pathogenic mutations (patients 1 and 2), or one pathogenic mutation and two SNPs (patient 3). Limiting dilution analysis was conducted using the Extreme Limiting Dilution Analysis software^16^, where the AML spike-in cell number was treated as ‘Dose’, and the number of replicates in which the leukemic fraction was detected was treated as ‘Response’. Output of the analysis provided an estimated sensitivity with an associated confidence interval.

### Plotting and Graphical Representation

All bar plots and scatter plots were generated using the ggplot2 package in R. The OncoPrint shown in Fig 2A was produced using the Complex Heatmap package in R. All heatmaps were generated using the pheatmap R package. The UMAP plots, density plots, and violin plots, in Fig 3 and Supplementary Fig 4 were generated using the Seurat R package. The radar plot displayed in Supplementary Fig 4 was produced with the fmsb package in R.

## Acknowledgement

This study was funded by the Center for Hematologic Malignancies at MSKCC and in part through the NIH/NCI Cancer Center Support Grant P30 CA008748. TR is supported by the American Society for Hematology (ASH). L.A.M. is supported by a National Cancer Institute grant (K99 CA252002-01A1). S.F.C. is supported by a Career Development Award from the NCI K08 CA241371-01A1. R.L.L. is supported by a Cycle for Survival Innovation Grant and National Cancer Institute R35 CA197594. WX is supported by Alex’s Lemonade Stand Foundation and the Runx1 Research Program, MSK’s Cycle for Survival’s Equinox Innovation Award in Rare Cancers, MSK Leukemia SPORE Career Enhancement Program and a National Cancer Institute grant (K08CA267058-01).

## Author contribution

W.X. selected, annotated and processed samples and prepared scDNA+protein libraries. T.R. performed computational analysis. R.L.B. and A.S. helped computational analysis. S.P. helped process normal bone marrow samples. Y.L. helped annotate bulk NGS sequencing data. Q.G., J.Z., X.T. and C.F. provided clinical samples. L.A.M. and S.F.C. helped scDNA+protein library preparation. A.L., A.G. and A.D. helped design the LOD study. M.R., R.L., and W.X. designed and supervised the entire study. T.R., R.L. and W.X. wrote the manuscript.

