## supplementary methods and figures for "Single cell genotypic and phenotypic analysis of measurable residual disease in acute myeloid leukemia"

*scMRD Computational Demultiplexing*

Deconvolution of multiplexed scMRD runs was reliant on the presence of germline SNPs. Suspected SNPs were verified via referencing the Ensembl SNP database through the BioMart R package and were tallied for non-missing genotyping information within the filtered NGT matrix. The top 10-20 SNPs with the lowest percentage of missing genotypes were selected for downstream analysis. K-means clustering was performed on SNP allele frequencies in a subset of cells with complete SNP genotypes, where the number of clusters for partitioning was set equal to the number of unique patient samples in a given multiplex. Doublet identification and exclusion was conducted by first evenly sampling cells from all clusters to form a pool of cells with equal representation of each cluster. Artificial doublets were then generated via sampling the cell pool two cells at random and averaging the SNP profiles until the proportion of artificial doublets approached 5-10% of the total number of cells in the dataset. Doublets were then merged with real cells and re-clustered to produce real and artificial cluster centers. The Euclidean distance was then measured between each real cell and 1) it’s respective cluster, 2) the artificial cluster center. The distribution of distances between 90-95% of cells to their respective cluster centers was used as a cutoff to exclude cells which were within this distance to the artificial cluster center. This process was repeated 10 times, with random replacement of NA values with allele frequencies of 0, 50, or 100, and cells were excluded if their distance was within the doublet gate in all replicates. After removing doublets and low-quality cells with high similarity to artificial doublets, the most common SNP profile was tallied for each cluster. To classify additional cells, a Hamming distance was calculated between all cells and each SNP profile, without penalizing SNPs with missing genotypes. Cells were assigned to clusters based on matching 80% of the SNP profile and being the maximum Hamming distance from every other cluster. For some multiplexed runs, slightly less stringent filters were applied to reduce the Hamming distance between clusters. After cell classification, each cluster was queried for pathogenic mutations detected by bulk NGS at the diagnosis, remission, and relapse (if applicable) timepoints, and the cell number per cluster was tallied.

*Single Cell Protein Analysis*

For each demultiplexed sample, single cell protein data was extracted from H5 files as raw counts. Each demultiplexed patient sample was analyzed independently for clonality of mutations and clone-specific immunophenotype. For samples with detected mutations, the protein count matrices were filtered for cells classified into high-confidence clones (>3 cells) and were used for subsequent aggregate analysis. Protein counts for each run were merged and converted to a Seurat object using the Seurat R package. The protein data was log-normalized, scaled, and centered on a by-run basis. Clone and mutation information was supplied as metadata and used for downstream aggregate analysis using functions within Seurat.

**Supplementary Tables**

See separate excel file

**Supplementary Figures**

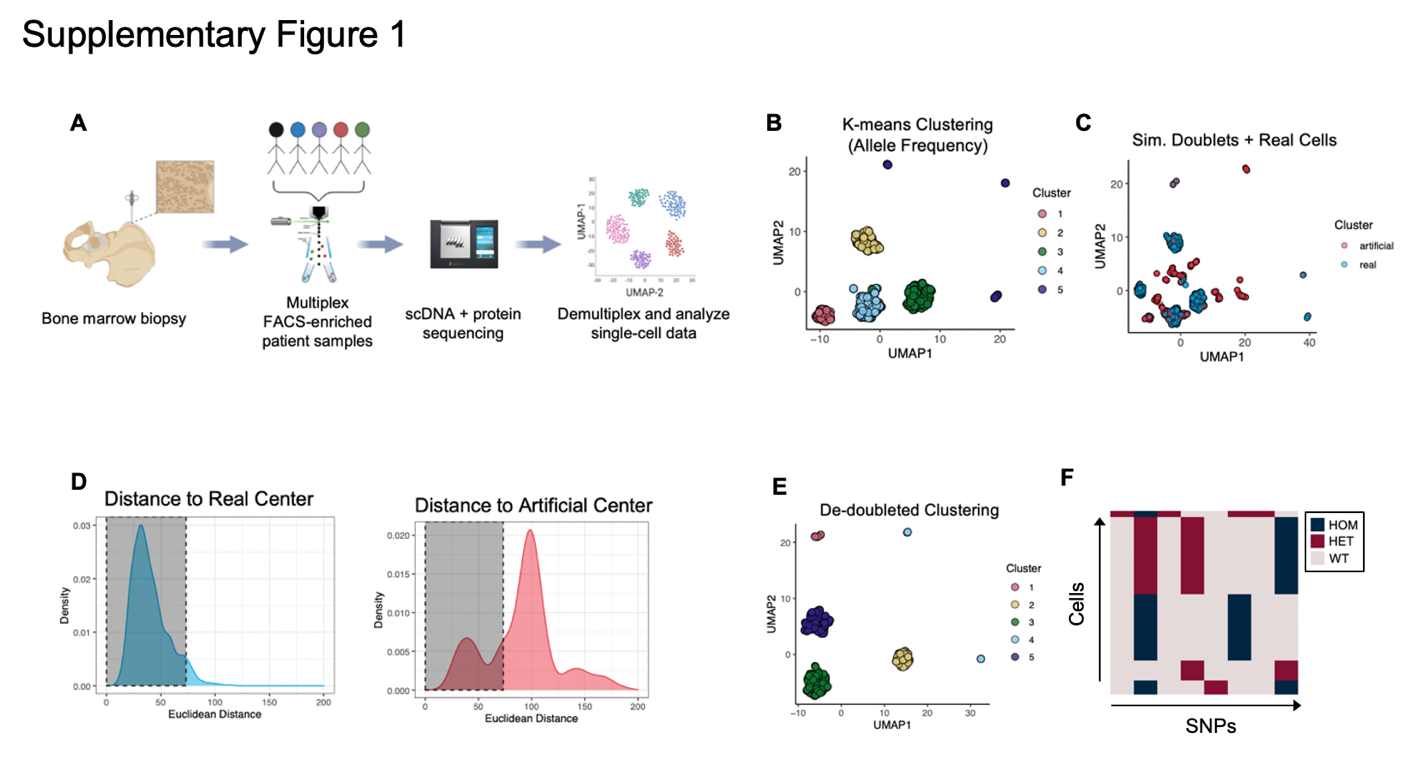

**Figure S1**. Workflow and computational demultiplexing of scMRD data. **A**. Schema of scMRD workflow (generated via BioRender). **B**. K-means clustering and UMAP analysis of SNP allele frequencies before doublet exclusion. **C**. UMAP plot showing the results of clustering real cells (blue) with artificial doublets (red). **D**. Distribution of Euclidean distances from real cells to their respective cluster centers (left) and to the artificial cluster center (right). **E**. K-means clustering and UMAP analysis of SNP allele frequencies after doublet exclusion. **F**. Heatmap showing private SNP genotypes in singlet clusters. Panels B-F show representative examples of the computational pipeline output.

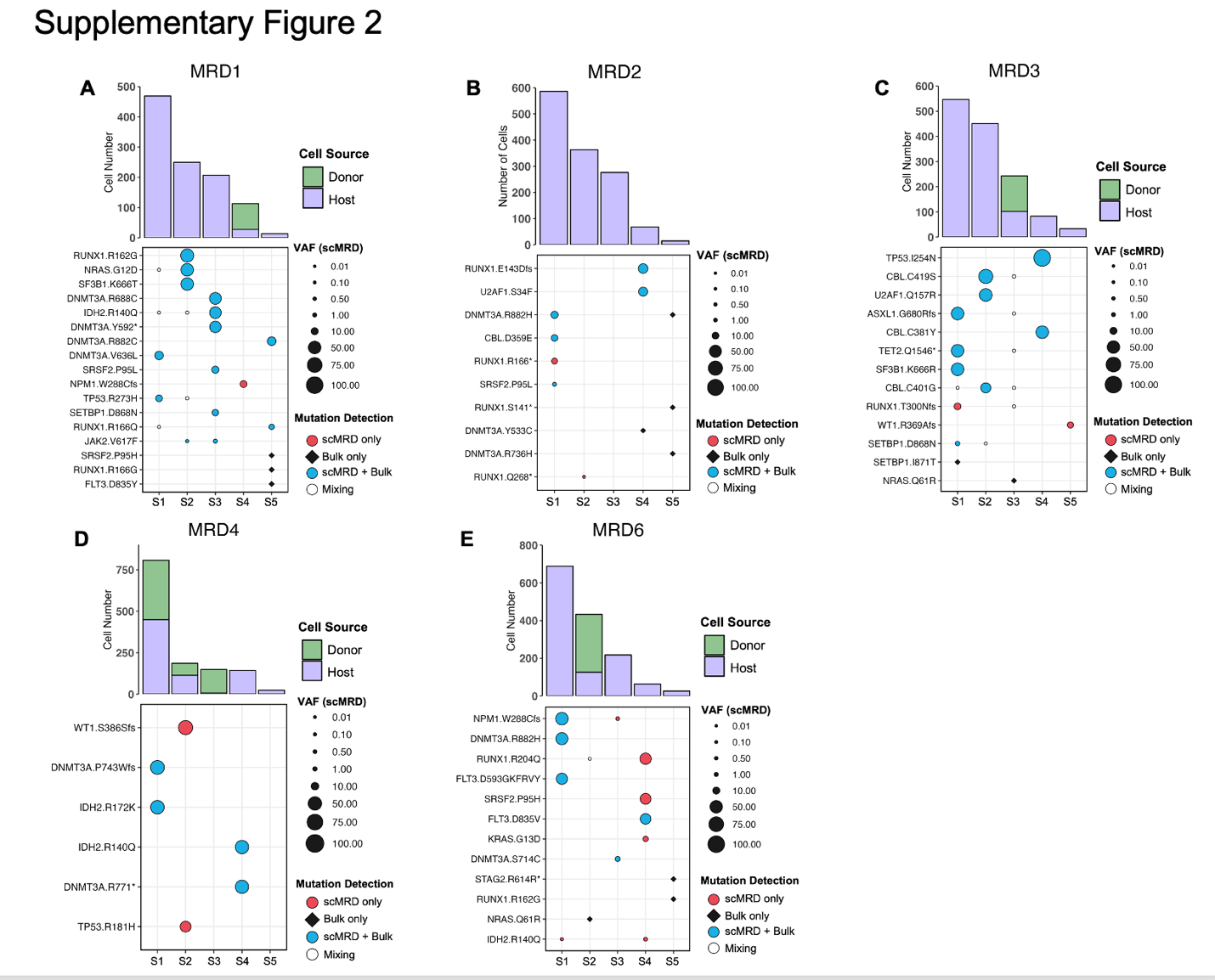

**Figure S2**. Deconvolution plots for scMRD runs. **A-E**. Recovered cell number per sample (top) and VAF of mutations detected by scMRD, bulk NGS, or both assays (bottom). Mixing represents mutations found in ≤2 cells that were likely misclassified by the demultiplexing pipeline.

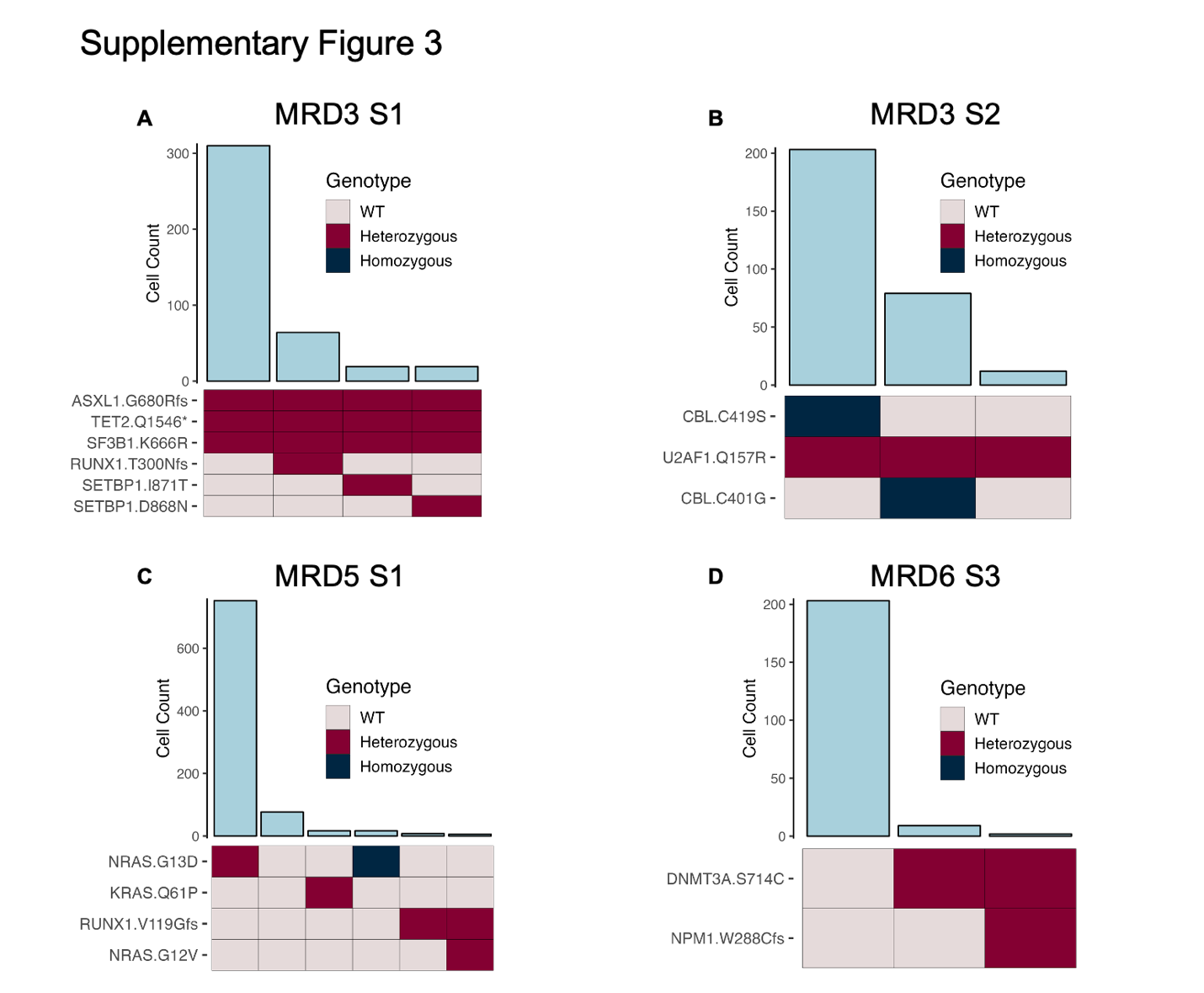

**Figure S3**. Representative clonographs of MRD samples. **A-D**. Columns represent individual clones identified in each sample, with cell count (top, bar plot) and zygosity of mutations present (bottom, heatmap).

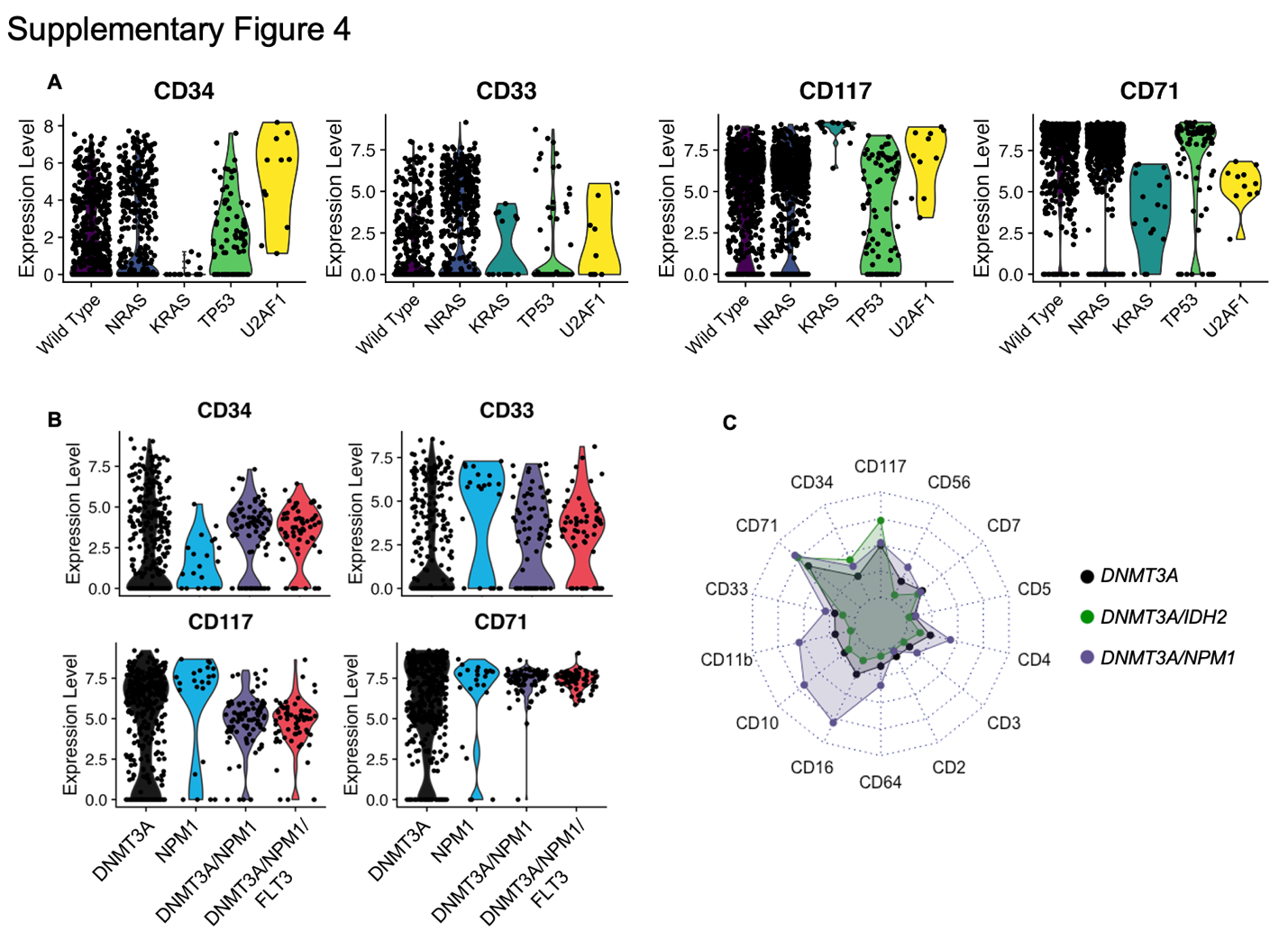

**Figure S4**. Analysis of protein sequencing data of MRD clones. **A**. Violin plots showing log- normalized differential surface marker expression of various MRD clones. **B**. Violin plots showing log-normalized differential surface marker expression of CH/preleukemic (*DNMT3A*) vs leukemic (*NPM1*, *DNMT3A/NPM1*, *DNMT3A/NPM1/FLT3ITD)* clones. **C**. Radar plot showing differential surface marker expression of CH/preleukemic (*DNMT3A*) vs leukemic (*DNMT3A/NPM1*, *DNMT3A/IDH2*) clones. Each marker is scaled relative to the maximum and minimum expression values for all cells with *DNMT3A*, *DNMT3A/NPM1*, or *DNMT3A/IDH2* mutations.

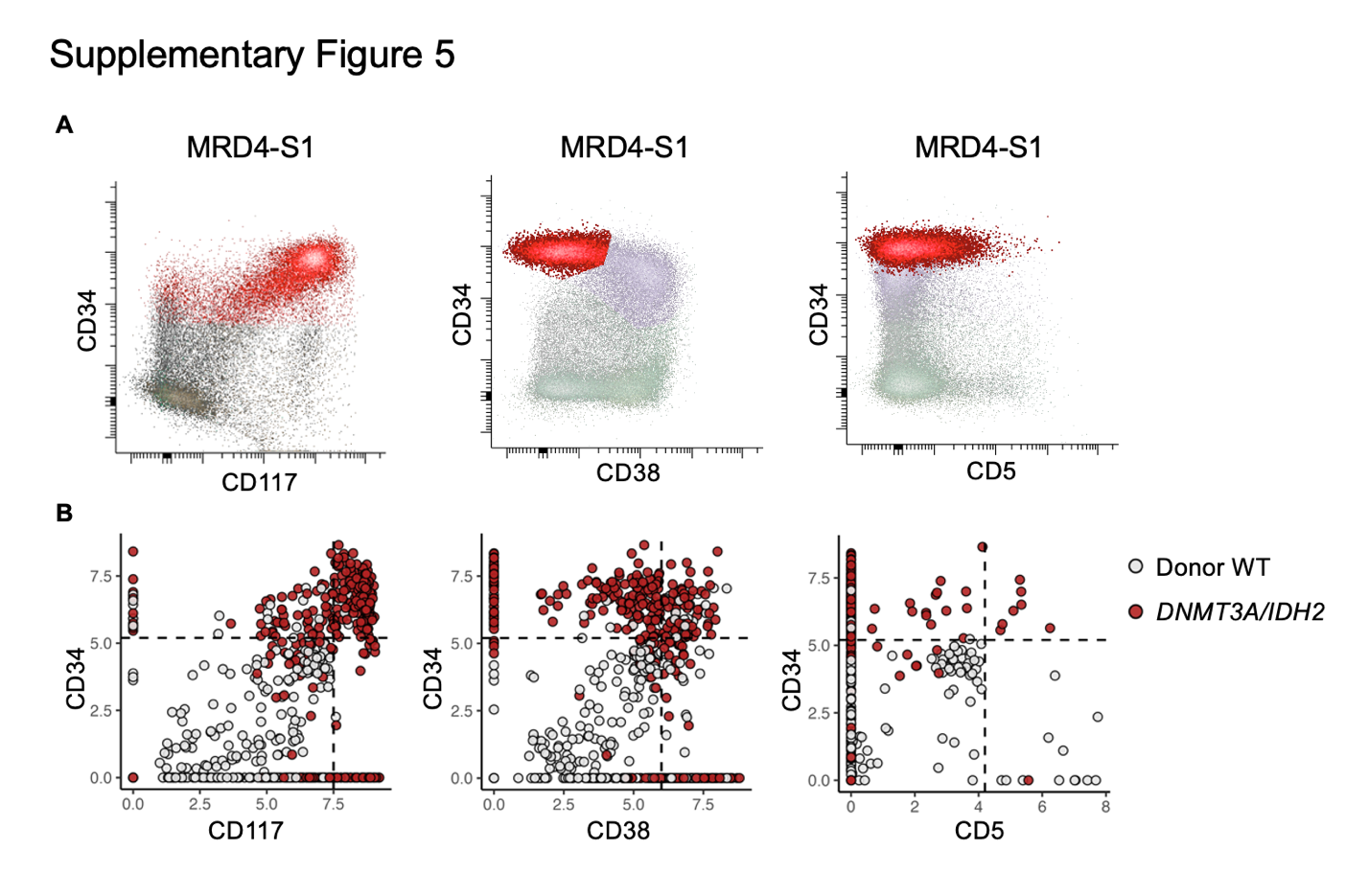

**Figure S5**. Concordance of immunophenotype between MFC and scMRD assay from a representative patient (MRD4-S1). **A**. Flow plots showing abnormal expression of bright CD117, dim to negative CD38 and partial CD5 on CD34 positive myeloblasts. **B**. scMRD data shows similar immunophenotype.
